# Fusion Speed of Biomolecular Condensates

**DOI:** 10.1101/2020.06.22.164897

**Authors:** Archishman Ghosh, Huan-Xiang Zhou

**Author notes:** Supporting information for this article is given via a link at the end of the document.

## Abstract

Biomolecular condensates formed through phase separation have a tendency to fuse. The speed with which fusion occurs is a direct indicator of condensate liquidity, which is key to both cellular functions and diseases. Using a dual-trap optical tweezers setup, we found the fusion speeds of four types of condensates to differ by two orders of magnitude. The order of fusion speed correlates with the fluorescence of Thioflavin T, which in turn reflects the macromolecular packing density inside condensates. Unstructured protein or polymer chains pack loosely and readily rearrange, leading to fast fusion. In contrast, structured protein domains pack more closely and have to break extensive contacts before rearrangement, corresponding to slower fusion. This molecular interpretation for disparate fusion speeds portends a unified understanding of the underlying physicochemical determinants.

**Entry for the Table of Contents:** The tendency of biomolecular condensates to fuse is key to cellular function and diseases. Using optical tweezers, fluorescence microscopy, and theoretical modeling, Ghosh and Zhou have begun to unravel the molecular origin for disparate fusion speeds among different biomolecular condensates. They found that fusion speed is dictated by macromolecular packing density inside condensates, which can be reported by ThT fluorescence.

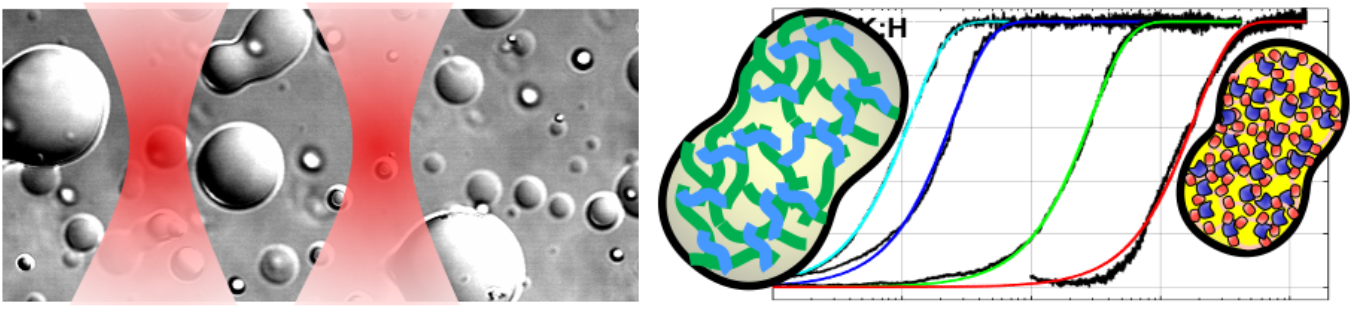

---

Biomolecular condensates form the hubs of many cellular functions such as transcription and stress response.^[1–2]^ These condensates include membraneless organelles such as nucleoli, P granules, and stress granules; typically appear as micro-sized droplets; and span material states from liquid-like to solid-like (or becoming solid-like over time, a process known as aging). In many cases, the liquid state is crucial for functionality and solidification is linked to neurodegeneration and other pathologies.^[3–4]^ Thus an ATP-dependent process actively maintains the internal fluidity of nucleoli,^[5]^ and RNA has been shown to increase the fluidity of LAF-1 protein droplets^[6]^ and proposed to prevent condensates formed by prion-like RNA-binding proteins from solidification.^[7]^

The speed of fusion between droplets is a direct indicator of liquidity: fast corresponds to a more liquid-like state whereas slow fusion a more solid-like state. A number of studies have reported fusion speeds, using either fluorescence microscopy^[5–6, 8–11]^ or optical tweezers (OT).^[3–4, 12–14]^ In the former case, one tracks the time change in the shape (e.g., aspect ratio or neck radius) of two fusing droplets. In the latter case, one tracks the forces that the fusing droplets exert as they move away from the optical traps. It is generally understood that fusion is accelerated by the interfacial surface tension (*γ*) of the droplets and retarded by viscosity (in particular, interior viscosity *η* when the bulk phase has much lower viscosity), and thus the fusion time (*τ*_*f*_) is proportional to *η* and inversely proportional to γ. Observed fusion times of related systems can differ by one to two orders of magnitude.^[5, 9, 11–12, 14]^

We are just starting to identify and understand the physicochemical determinants of condensate fusion speed. Several studies have highlighted the role of the strength of intermolecular local interactions. For example, disease-linked mutations that increase the propensity for protein aggregation, presumably by strengthening local interactions, also slow down fusion.^[3–4]^ Similarly, the fusion times of binary droplets formed by arginine- or lysine-containing peptides with purine or pyrimidine homopolymeric RNAs follow the order of the strengths of cation-pi interactions.^[11, 14]^ Another determinant is molecular flexibility, as indicated by a 350-fold increase in fusion time when a subset of glycine residues in the prion-like domain of FUS was mutated into alanine’s.^[12]^ Macromolecular crowding has been reported to slow down fusion, although different mechanisms may exist.^[10, 13]^ There was also a suggestion that electrostatic interactions confer liquid-like properties whereas hydrophobic interactions confer solid-like properties,^[15]^ but its generality was questioned.^[16]^ Complicating the matter, some factors, e.g., interaction strength, may affect both surface tension and viscosity in the same direction, leaving the net effect on fusion time uncertain. For example, Increasing salt concentration, by weakening electrostatic interactions, reduced viscosity inside LAF-1 droplets,^[6]^ but by the same token, surface tension is also expected to be reduced. The effect of increasing salt concentration on *τ*_*f*_ was not reported, but would depend on whether the salt has a greater effect on surface tension or viscosity. Droplets separately formed by two major components of nucleoli, nucleophosmin and fibrillarin, provide a clear demonstration of the tussle between surface tension and viscosity.^[9]^ Compared to nucleophosmin droplets, fibrillarin droplets have a higher surface tension, which would make them fuse faster, but in fact their fusion time is over 100-fold longer, implicating viscosity as the overriding factor in this case. Lastly, we note our recent proposal of the structural compactness of component macromolecules as a determinant of material properties.^[16]^

Here we present quantitative analysis of OT-tracked fusion progress curves for a variety of droplets (Figure 1) to gain better understanding of the physicochemical determinants of fusion speed. To provide a theoretical basis for the quantitative analysis, we model the droplet phase as an incompressible, viscous fluid, and the fusion process as a Stokes flow (see Supporting Information, Section 5). The reduction in the edge-to-edge distance, *l*, fits well to a stretched exponential,

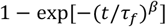

with *β* = 1.5 and *τ*_*f*_ = 1.56(*η*/*γ*)*a*, where *a* denotes the radius of the droplets (Figure S1). We prepared the two fusing droplets in equal size (Supporting Information, Section 2). The difference between the forces detected by the two optical traps is proportional to the reduction in *l* (Supporting Information, Section 6; Figure S2), and thus fits also to the above stretched exponential (Figure 1b). The fusion times (*τ*_*f*_) of four types of droplets, all with similar sizes, span over two orders of magnitude.

**Figure 1.**
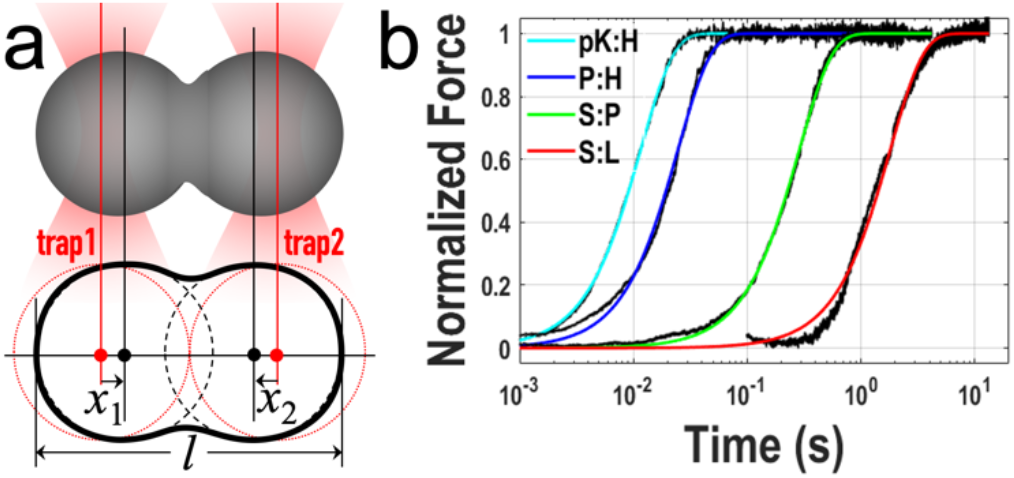
Droplet fusion process tracked by optical tweezers. a) Illustration of the fusion of two droplets under optical trapping. b) Force traces of 4 different types of droplets, with similar initial sizes (radii ranging from 1.9 to 2.9 μm).

Three of the four types of droplets were formed by binary mixtures, P:H, S:P, and S:L, of four macromolecular species: pentameric constructs of SH3 domains (S; acidic) and proline-rich motifs (P; basic), single-domain protein lysozyme (L; basic), and polymer heparin (H; acidic). L and H were identified as representatives of suppressors and promoters, respectively, of S:P phase separation.^[17]^ A fourth binary, H:L, formed network-like precipitates^[16]^ and hence were unsuited for fusion study by OT. We chose another polymeric promoter, polylysine (pK),^[17]^ whose mixtures with H also formed network-like precipitates at low salt but turned into droplets at 0.8 M KCl (Figure S3), similar to a previous study.^[18]^ The pK:H droplets have the shortest fusion time. In comparison, P:H, S:P, and S:L droplets fuse at speeds approximately 2-, 20-, and 100-fold lower.

The Stokes model predicts a proportional relation between fusion time and droplet size. We measured fusion times for droplets with radii ranging from 0.8 to 8 μm, and they indeed follow a proportional relation (Figure 2a), as also reported in some previous studies.^[5–6, 8–9]^ The slope *S* ≡ *τ*_*f*_/*a*, equal to 1.56*η*/*γ* according to the Stokes model, is 6.7, 13.1, 128.0, and 591.2 ms/μm, respectively, for pK:H, P:H, S:P and S:L droplets. In previous work, we mixed the viscosity-sensitive dye, thioflavin T (ThT), into P:H, S:P and S:L droplets, and found background-level, weak, and strong fluorescence, respectively, in fluorescence these three types of droplets.^[16]^ This ranking in fluorescence intensity is identical to the ranking in fusion time, hinting that viscosity dominates over surface tension in controlling fusion speed. Indeed, *S* exhibits a linear correlation with the ratio of the ThT fluorescence intensity in the droplet phase to that in the bulk phase (Figure 2b, c).

**Figure 2.**
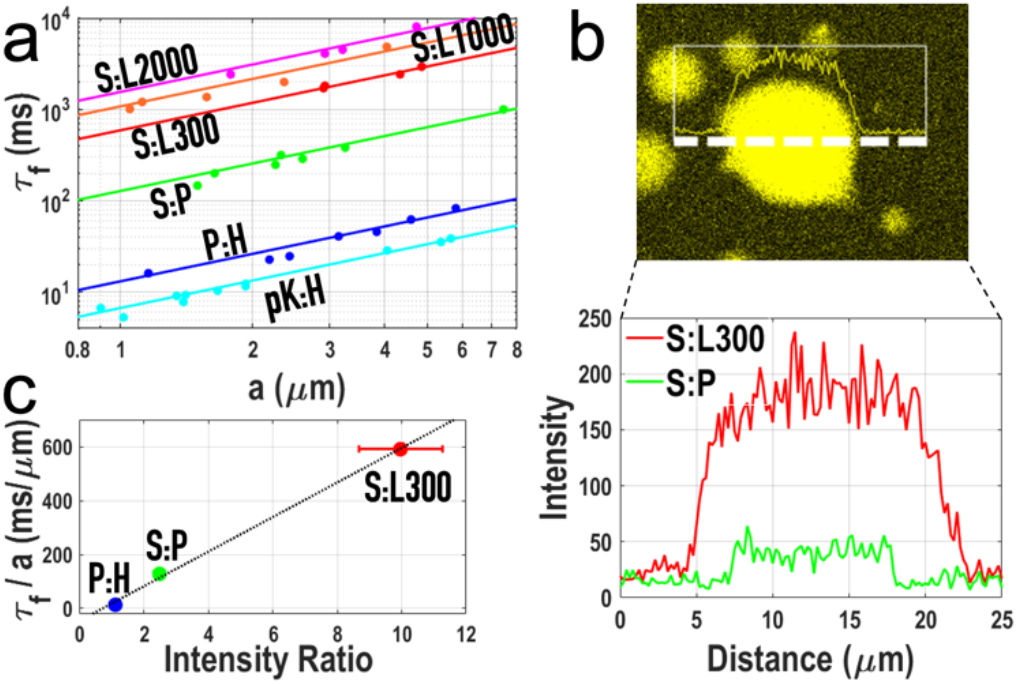
Correlation of fusion time with ThT fluorescence intensity. a) Proportional relation between fusion time (*τ*_*f*_) and droplet radius (*a*). Errors bars are smaller than the size of the symbols. b) A line scan of ThT fluorescence across an S:L droplet, and the intensity profiles for this S:L droplet and for an S:P droplet, illustrating the contrast in ThT fluorescence between these droplets. c) Linear relation between *τ*_*f*_/*a* and the ratio of ThT fluorescence intensities inside and outside droplets. Some errors are not visible because they are smaller than the size of the symbols.

Further support for viscosity as the dominant factor for fusion speed came from S:L droplets prepared with higher L concentrations. Upon increasing L from 300 μM to 1000 and 2000 μM, *S* increases from 591.2 ms/μm to 1080.5 and 1554 ms/μm, respectively (Figures S4a and 2a). The increase in *S* comes as the L concentration inside droplets also increases (Figure S4b). Interestingly, S:L droplets prepared at 486 μM of L collapse on a coverslip but those at 1736 μM of L stand as spherical, indicating an increase in surface tension at the higher L. Thus, with increasing L, viscosity must increase at a higher rate than surface tension. In fact, the viscosity of L solutions is a rapidly increasing function of concentration,^[19]^ and ThT fluorescence intensities increase commensurately (Figure S5).

According to the Stokes model, a universal function, 1 − exp[−(*t*/*τ*_*f*_)^*β*^, fits all OT-detected force progress curves. If time is scaled by the respective fusion times, then all progress curves should collapse to a master curve. Figure 3 shows that this is indeed the case, providing strong validation of the Stokes model.

**Figure 3.**
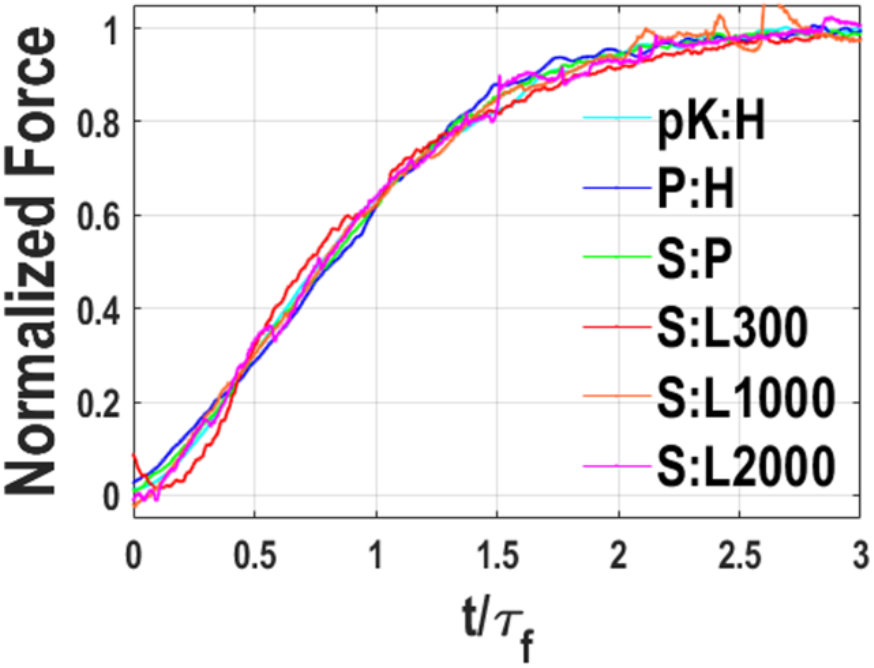
Collapse of fusion progress curves into a single master curve. Progress curves were smoothed over periods ranging from 0.64 to 32 ms in duration.

The correlation between fusion time and ThT fluorescence provides a starting point for seeking the molecular origin of the wide disparity in fusion speed between the different types of droplets studied here. Previously we have interpreted ThT fluorescence as reflecting macromolecular packing density, which in turn is related to the structural compactness of component macromolecules and to their interactions.^[16]^ Let us consider the two extremes, i.e., S:L and pK:H droplets, in the *τ*_*f*_ spectrum. S comprises five flexibly-linked, structured, SH3 domains. In S:L droplets, each single-domain L molecule is surrounded by SH3 domains and linkers from multiple S molecules, likely including at least one pair of L and SH3 domains with extensive contacts and leading to close packing (Figure 4a). In contrast, pK and H are unstructured polymer chains. Intermolecular interactions in pK:H droplets largely involve physical crosslinks between the chain molecules, corresponding to loose packing (Figure 4b). Fusion requires rearrangement of the macromolecular matrices inside two droplets, and can be rate-limited by the breakup of inter-domain contacts (as in S:L droplets) or inter-chain crosslinks (as in pK:H droplets). Breaking the extensive contacts between two structured domains takes more energy and hence occurs more slowly than breaking a physical crosslink between two unstructured chains, a distinction noted previously.^[20]^ Rearrangement of unstructured chains is further aided by their molecular flexibility. These differences explain the much slower fusion of S:L droplets relative to pK:H droplets.

**Figure 4.**
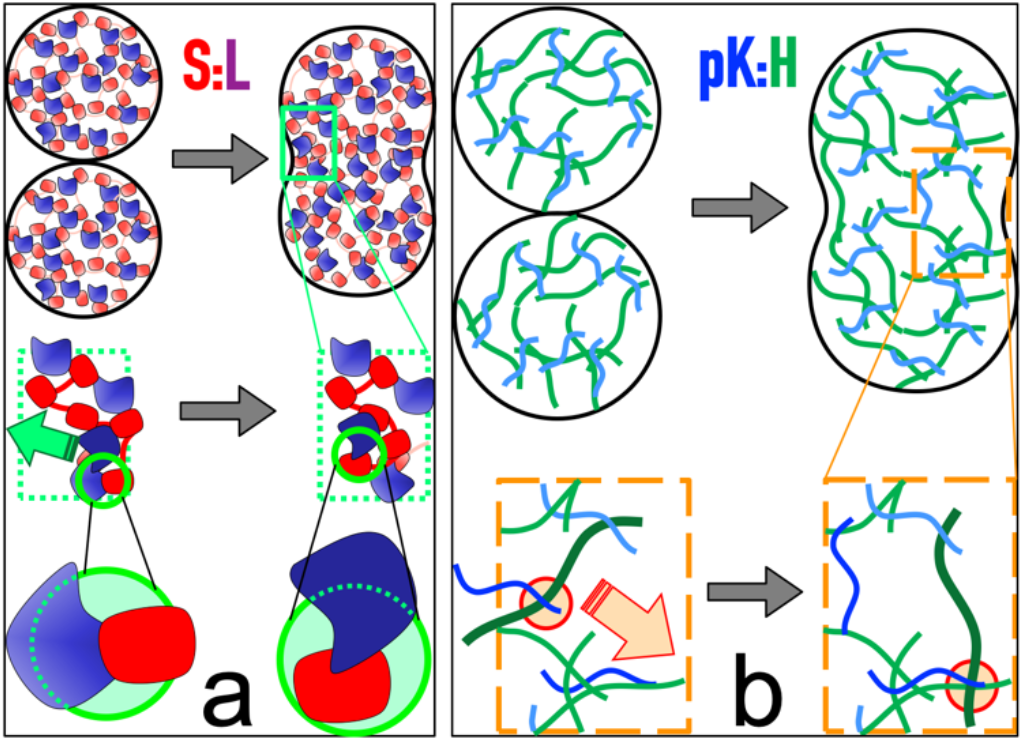
Differences between S:L and pK:H droplets in packing and in contact breaking during fusion. a & b) In S:L (pK:H) droplets, macromolecules are closely (loosely) packed; fusion is rate-limited by breaking old inter-domain contacts (inter-chain physical crosslinks) before forming new ones.

P and H are also unstructured polymer chains and hence P:H and pK:H droplets have similar fusion speeds. The 2-fold higher fusion speed of pK:H droplets may result from the higher KCl concentration that we had to work with, leading to weakened electrostatic attraction. S:P droplets present an intermediate situation, where intermolecular interactions consist of both domain-chain contacts (between SH3 in S and proline-rich motif in P) and interchain crosslinks (involving linkers in S). The resulting packing density and fusion speed are also intermediate between those of P:H and S:L droplets. Fusion speed and ThT fluorescence intensity are correlated because they are both dictated by macromolecular packing density inside droplets.

In summary, our data support two conclusions. First, structural compactness of component macromolecules is a new determinant of fusion speed. Second, packing density, which can be reported by ThT fluorescence, integrates this and other determinants into a single fusion-speed predictor. The first conclusion explains why deleting a structured domain in fibrillarin led to a dramatic, 80-fold speedup in fusion,^[9]^ whereas the second conclusion favors increased component concentration inside droplets, through displacement by crowders in the bulk phase,^[17, 21]^ as a mechanism for the slowdown of fusion by macromolecular crowding.^[13]^ We anticipate that these conclusions will stimulate further investigations into the physicochemical determinants of condensate fusion speed, leading to a unified understanding.

## Acknowledgements

This work was supported by National Institutes of Health Grant GM118091.

## References

[1] W. K. Cho, J. H. Spille, M. Hecht, C. Lee, C. Li, V. Grube, Cisse, II, Science 2018, 361, 412–415.

[2] S. Kroschwald, M. C. Munder, S. Maharana, T. M. Franzmann, D. Richter, M. Ruer, A. A. Hyman, S. Alberti, Cell Rep 2018, 23, 3327–3339.

[3] A. Patel, H. O. Lee, L. Jawerth, S. Maharana, M. Jahnel, M. Y. Hein, S. Stoynov, J. Mahamid, S. Saha, T. M. Franzmann, A. Pozniakovski, I. Poser, N. Maghelli, L. A. Royer, M. Weigert, E. W. Myers, S. Grill, D. Drechsel, A. A. Hyman, S. Alberti, Cell 2015, 162, 1066–1077.

[4] X. Gui, F. Luo, Y. Li, H. Zhou, Z. Qin, Z. Liu, J. Gu, M. Xie, K. Zhao, B. Dai, W. S. Shin, J. He, L. He, L. Jiang, M. Zhao, B. Sun, X. Li, C. Liu, D. Li, Nat Commun 2019, 10, 2006.

[5] C. P. Brangwynne, T. J. Mitchison, A. A. Hyman, Proc Natl Acad Sci U S A 2011, 108, 4334–4339.

[6] S. Elbaum-Garfinkle, Y. Kim, K. Szczepaniak, C. C. H. Chen, C. R. Eckmann, S. Myong, C. P. Brangwynne, Proc Natl Acad Sci U S A 2015, 112, 7189–7194.

[7] S. Maharana, J. Wang, D. K. Papadopoulos, D. Richter, A. Pozniakovsky, I. Poser, M. Bickle, S. Rizk, J. Guillen-Boixet, T. M. Franzmann, M. Jahnel, L. Marrone, Y. T. Chang, J. Sterneckert, P. Tomancak, A. A. Hyman, S. Alberti, Science 2018, 360, 918–921.

[8] C. P. Brangwynne, C. R. Eckmann, D. S. Courson, A. Rybarska, C. Hoege, J. Gharakhani, F. Jülicher, A. A. Hyman, Science 2009, 324, 1729–1732.

[9] M. Feric, N. Vaidya, T. S. Harmon, D. M. Mitrea, L. Zhu, T. M. Richardson, R. W. Kriwacki, R. V. Pappu, C. P. Brangwynne, Cell 2016, 165, 1686–1697.

[10] C. M. Caragine, S. C. Haley, A. Zidovska, Phys Rev Lett 2018, 121, 148101.

[11] S. Boeynaems, A. S. Holehouse, V. Weinhardt, D. Kovacs, J. Van Lindt, C. Larabell, L. V. D. Bosch, R. Das, P. S. Tompa, R. V. Pappu, A. D. Gitler, Proc Natl Acad Sci U S A 2019, 116, 7889–7898.

[12] J. Wang, J. M. Choi, A. S. Holehouse, H. O. Lee, X. Zhang, M. Jahnel, S. Maharana, R. Lemaitre, A. Pozniakovsky, D. Drechsel, I. Poser, R. V. Pappu, S. Alberti, A. A. Hyman, Cell 2018, 174, 688–699.

[13] T. Kaur, I. Alshareedah, W. Wang, J. Ngo, M. M. Moosa, P. R. Banerjee, Biomolecules 2019, 9.

[14] I. Alshareedah, T. Kaur, J. Ngo, H. Seppala, L. A. D. Kounatse, W. Wang, M. M. Moosa, P. R. Banerjee, Journal of the American Chemical Society 2019.

[15] S. C. Weber, Curr Opin Cell Biol 2017, 46, 62–71.

[16] A. Ghosh, X. Zhang, H. X. Zhou, J Am Chem Soc 2020, in press.

[17] A. Ghosh, K. Mazarakos, H. X. Zhou, Proc Natl Acad Sci U S A 2019, 116, 19474–19483.

[18] H. Zhou, S. Zhong, Z. Song, L. Zuo, Z. Qi, L. J. Qu, L. Lai, Angew Chem Int Ed Engl 2019, 58, 4858–4862.

[19] P. D. Godfrin, S. D. Hudson, K. Hong, L. Porcar, P. Falus, N. J. Wagner, Y. Liu, Phys Rev Lett 2015, 115, 228302.

[20] H. X. Zhou, Trends Biochem Sci 2012, 37, 43–48.

[21] V. Nguemaha, H. X. Zhou, Sci Rep 2018, 8, 6728.

