## Supplementary Text and Figures for "Fusion Speed of Biomolecular Condensates"

**Experimental Procedures**

*1. Sample preparation*. Pentameric constructs of SH3 domains (S) and proline-rich motifs (P) were expressed and purified as described previously.[1] The sources for lysozyme (L), FITC-lysozyme, heparin (H) , polylysine (pK) , Thioflavin-T (ThT), and Ficoll70 were the same as reported previously.[1-2] All experiments were conducted in a buffer containing 10 mM imidazole pH 7, 0.01% $NaN_3$, and KCl at 0.15 M or an otherwise indicated concentration. The droplets for fusion experiments were prepared at the following concentrations in μM: pK:H = 50:50; P:H = 40:40; S:P = 40:40; and S:L = 20:300 or otherwise with L at 1000 or 2000. These samples also contained 50 g/L of Ficoll70 to aid in the trapping of droplets by slowing down their falling under gravity. For pK:H droplets only, the KCl concentration was 1.0 M.

*2. Tracking of droplet fusion by optical tweezers (OT)*. Droplet fusion was tracked on a LUMICKS C-Trap™ dual-trap OT instrument. 7-8 μL of a freshly prepared, droplet-forming binary mixture was loaded into a custom-made sample chamber, consisting of a coverslip (VWR cat# 48366-067) attached to the center of a slide (VWR cat# 48300-026) via two strips of double-sided tape (Staples cat# 378988) spaced 2-3 mm apart. The laser of the C-Trap™ (1064 nm wavelength) was tuned to low power, just enough to trap droplets on the basis of their difference in refractive index from the bulk phase. Two droplets were separately trapped and if necessary, allowed to scavenge nearby droplets until they reached equal size. The two droplets were then held in separation until all nearby droplets in the field had fallen under gravity. The droplet in trap 1 was next moved close to the droplet in trap 2. A snapshot of the two still separated droplets were taken, and the video recording was turned on (with a 110 μm × 88 μm field of view at a 1280 × 1024 pixel resolution). Care was taken to align the traps along a horizontal axis ($x$ axis). The trap-1 droplet was finally moved in steps of 10 or 20 nm until it came into contact with the trap-2 droplet to start fusion, which continued without any disturbance. Following completion of fusion, the force profile was saved, and video recording was terminated. The force sampling rate was 78.125 kHz (corresponding to a time resolution of 12.8 μs) and the video frame rate was 25 per s.

*3. Analysis of force progress curves*. The progress curves for the $x$ components of the forces detected by the two traps mirrored each other (trap 1 negative whereas trap 2 positive), albeit trap 1 was more prone to perturbations because of its involvement in pre-fusion maneuver. We thus chose to analyze only the force curve from trap 2, by fitting it to a stretched exponential function in MATLAB. The function had the following form:

$$Force = A + B\left\{1 - e^{-[(t-t_0)/\tau_f]^{\beta}}\right\} H(t - t_0)$$

where $A$ and $A + B$ are the pre-fusion baseline and post-fusion plateau, respectively; $t_0$ denotes the start time of the fusion process; a Heaviside step function $H(t - t_0)$ was introduced to ensure that the stretched exponential applied only from time $t_0$ on; $\tau_f$ is a parameter that provides a measure for the fusion time; and $\beta$ was fixed to 1.5 (see below).

Our interest was the time course of the fusion process, and hence there was no need for calibrating the magnitudes of the forces detected by the traps. We did invoke the usual assumption of a proportional relation between force and displacement from the trap center, in order to justify the use of the stretched exponential for fitting the force curve (see below). We note that Wang et al.[3] have fitted OT-detected force progress curves to a stretched exponential, without justification. The interquartile range of their $\beta$ values were 1.30 to 2.05. We found it more useful to fix $\beta$ for comparing fusion speeds of different droplets.

*4. Image processing for droplet size measurements*. The radius of pre-fusion droplets was measured on the image that was saved just before fusion started. The image was opened in imageJ, and a circle was drawn to match the circumference of each droplet. The areas for both matching circles were measured in number of pixels and averaged. From this average area, the radius was calculated in μm, using the conversion factor 1 pixel = 0.0866 μm.

The time trace of the edge-to-edge distance was generated in MATLAB from the video recording of the fusing droplets. The video was first opened in imageJ, cropped, contrast-enhanced, and exported as a time sequence of image files in tiff. The MATLAB code then worked on each tiff file by image recognition, specifically by fitting the fusing droplets into a rectangular bounding box. The width of this box was reported as the edge-to-edge distance (sample output shown as green dots in Figure S2). In addition, a scale bar with length equal to the edge-to-edge distance was added to the image (sample output presented in the top row in Figure S2), and the time sequence of such images exported as a video (see Supporting Information Movie 1).

*5. Analysis of the numerical solution to the Stokes model for droplet fusion*. Following Martinez-Herrera and Derby,[4] we model the fusion between two droplets by the Stokes equations for an incompressible, viscous fluid:

$$\eta \nabla^2 \boldsymbol{v} - \nabla p = 0$$

$$\nabla \cdot \boldsymbol{v} = 0$$

where $\boldsymbol{v}$, $p$, and $\eta$ denote the velocity field, pressure field, and viscosity inside the droplets. The surrounding bulk phase is assumed not to affect the fusion process, except through the surface tension, $\gamma$, of the droplets. Martinez-Herrera and Derby numerically solved the Stokes equations for two fusing droplets. We found their progress curve for the edge-to-edge distance $l$ (Figure S1a) to fit well with a stretched exponential (Figure S1b):

$$l_0 - l = (l_0 - l_\infty)[1 - e^{-(t/\tau_f)^\beta}]$$

where $l_0$ and $l_\infty$ are the values of $l$ at time $t = 0$ and $\infty$, respectively, and $\beta$ is a parameter indicating the deviation from the usual exponential function. The best fit yielded $\beta = 1.43$ for two equal-sized droplets and this parameter decreased slightly for different-sized droplets. However, there is a strong correlation between $\beta$ and $\tau_f$. For simplicity, we fixed $\beta$ at 1.5. Then the value for $\tau_f$ is

$$\tau_f = 1.56(\eta/\gamma)a$$

Note that $\tau_f$ is proportional to $\eta$ and inversely proportional to $\gamma$, consistently with the expectation that viscosity slows down fusion whereas surface tension accelerates fusion. Moreover, the fusion time is proportional to the droplet radius $a$. The last relation served as an important check on the applicability of the Stokes model to our droplets (Figure 2a).

*6. Relation between edge-to-edge distance and force measured by optical traps*. As two droplets fuse, they move away from the centers of the dual traps. The traps thus each detect a force that is proportional to the displacement $x_i$, $i = 1$ or 2 (Figure 1a). Let the center of the first trap be the origin of the $x$ axis. The coordinate of the left edge of the first droplet is $l_1 = x_1 - a$; the coordinate of the right edge of the second droplet is $l_2 = 2a + x_2 + a$. The edge-to-edge distance is then $l = l_2 - l_1 = l_0 + x_2 - x_1$. This relation holds as a good approximation until near the end of the fusion process, when the fusing droplets can no longer be distinguished from each other. Consequently, for the reduction in $l$, we have $l_0 - l = x_1 - x_2$. Now $x_1 - x_2$ is proportional to the force difference between the two traps. So finally the force difference is proportional to $l_0 - l$, and can likewise be fitted to a stretched exponential. Because of the symmetry of the problem, the two forces should have opposite signs but equal magnitudes.

*7. Microscopy*. ThT fluorescence images of P:H, S:P, and S:L droplets and of lysozyme solutions were reported previously,[2] and analyzed here using the Zen Blue 2.5 software. Fluorescence intensities were measured over multiple regions of interest, drawn both inside and outside droplets. The intensity ratio was calculated after subtracting a baseline level (see next paragraph) from the intensities both inside and outside droplets (Figure 2c). For visual representation, a line scan was taken across a droplet, and the intensity profile along the line was exported for plotting (Figure 2b).

For each lysozyme solution, the entire field of view was treated as the region of interest. The dependence of fluorescence intensity on lysozyme concentration was fitted to a parabolic function (Figure S5), and the value extrapolated to 0 lysozyme concentration was used as the baseline level.

Confocal images of droplet-forming S:L mixtures (Figure S4b) were acquired on a Zeiss LSM 710 with a C-Apochromat 40× water objective. The samples were labeled with 2.5 μM FITC-lysozyme. The excitation wavelength was 488 nm and the detection wavelength range was 494-589 nm.

Images for pK:H condensates at different KCl concentrations (Figure S3) were acquired on an Olympus BX53 widefield microscope with a 40× air objective.

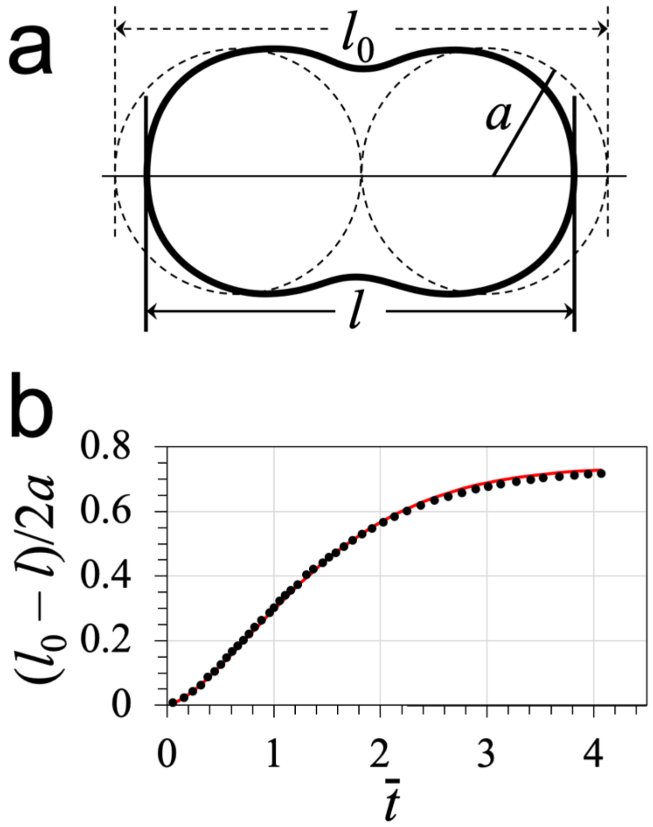


**Figure S1.** Solution of the Stokes model. a) Two equal-sized fusing droplets, with the edge-to-edge distance $l$ decreasing over time. The radius of the droplets is $a$. At time 0, the two droplets are tangent to each other, and the edge-to-edge distance $l_0$ equals $4a$. b) The dependence of the scaled reduction in edge-to-edge distance, $(l_0 - l)/2a$, on the dimensionless time, $\bar{t} \equiv (\gamma/\eta a)t$. Dots represent the numerical solution of Herrera· and Derby;[4] curve is the stretched exponential fit with $\beta$ = 1.5 and $(\gamma/\eta a)\tau_f$ = 1.56.

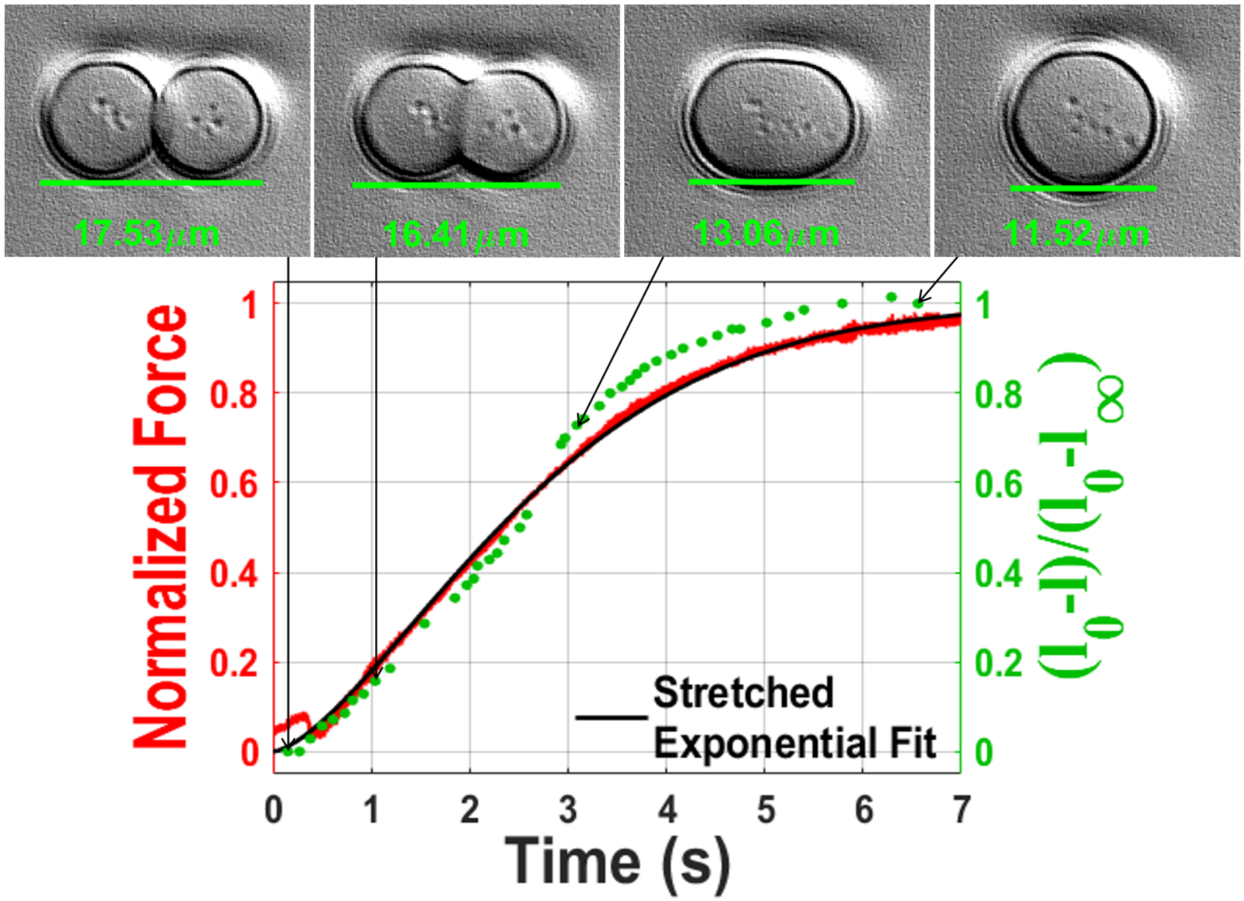


**Figure S2.** Comparison of the time traces of the OT-detected force (red) and the edge-to-edge distance from image processing (green), for two S:L droplets (at 20:300 μM). The top row presents four snapshots of the fusing droplets at different times; see Supporting Information Movie 1 for the entire fusion process.

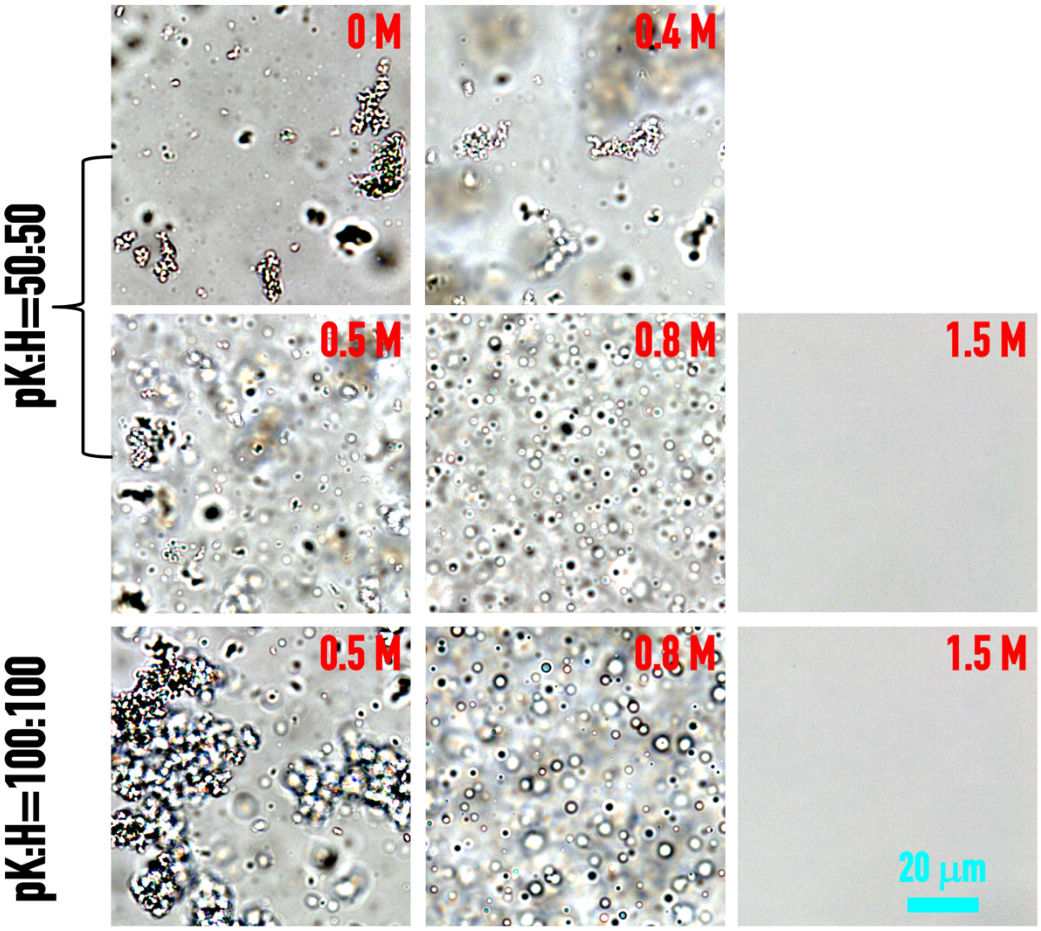


**Figure S3.** Condensates formed by pK:H mixtures at two indicated concentrations (in μM). Network-like precipitates were observed at KCl up to 0.5 M. Only droplets were observed at 0.8 M KCl. All condensates were dissolved at 1.5 M KCl.

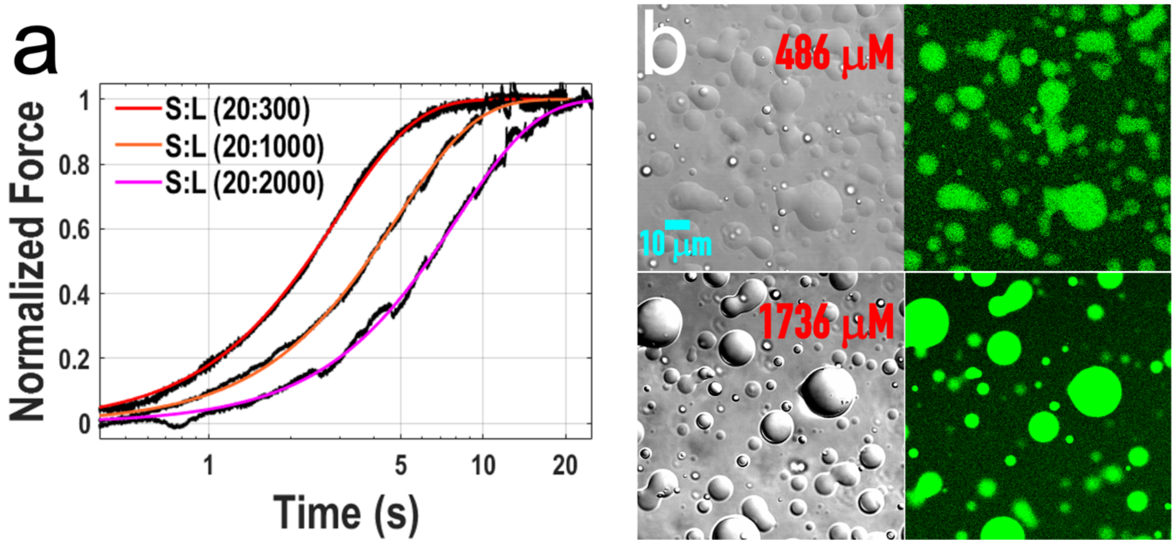


**Figure S4.** a) Representative force traces for the fusion of S:L droplets at three L concentrations (in μM). b) Brightfield and fluorescence images of S:L droplets at two L concentrations. S concentration was kept at 20 μM.

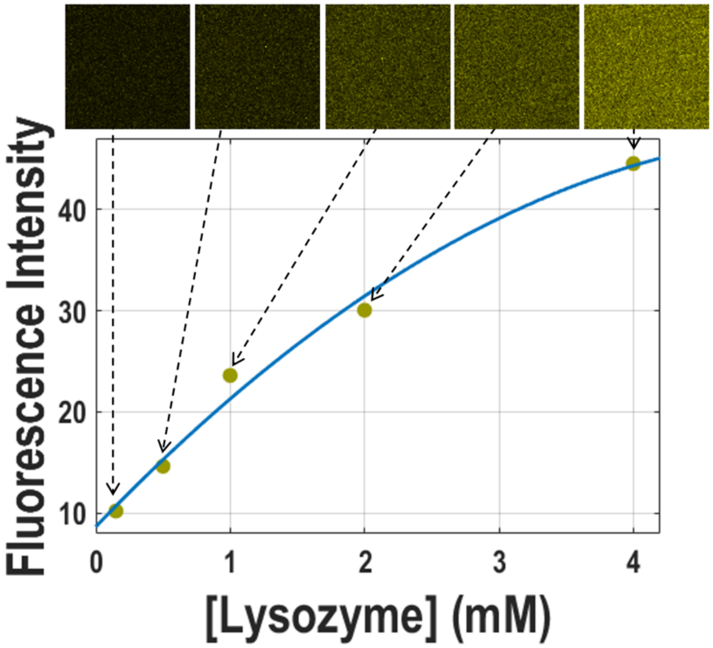


**Figure S5.** ThT fluorescence intensities in lysozyme solutions. Dots are intensities in arbitrary units; curve is a parabolic fit. The top row shows fluorescence images of ThT mixed into the lysozyme solutions; these images have appeared previously.[2]

### References


[1] A. Ghosh, K. Mazarakos, H. X. Zhou, *Proc Natl Acad Sci U S A* **2019**, *116*, 19474-19483.

[2] A. Ghosh, X. Zhang, H. X. Zhou, *J Am Chem Soc* **2020**, in press.

[3] J. Wang, J. M. Choi, A. S. Holehouse, H. O. Lee, X. Zhang, M. Jahnel, S. Maharana, R. Lemaitre, A. Pozniakovsky, D. Drechsel, I. Poser, R. V. Pappu, S. Alberti, A. A. Hyman, *Cell* **2018**, *174*, 688-699.

[4] J. I. Martínez-Herrera, J. J. Derby, *Journal of the American Ceramic Society* **1995**, *78*, 645-649.
